# Input dependent modulation of olfactory bulb activity by GABAergic basal forebrain projections

**DOI:** 10.1101/2020.03.29.014191

**Authors:** Erik Böhm, Daniela Brunert, Markus Rothermel

## Abstract

Basal forebrain modulation of central circuits is associated with active sensation, attention and learning. While cholinergic modulations have been studied extensively the effect of non-cholinergic basal forebrain subpopulations on sensory processing remains largely unclear. Here, we directly compare optogenetic manipulation effects of two major basal forebrain subpopulations on principal neuron activity in an early sensory processing area, i.e. mitral/tufted cells (MTCs) in the olfactory bulb. In contrast to cholinergic projections, which consistently increased MTC firing, activation of GABAergic fibers from basal forebrain to the olfactory bulb lead to differential modulation effects: while spontaneous MTC activity is mainly inhibited, odor evoked firing is predominantly enhanced. Moreover, sniff triggered averages revealed an enhancement of maximal sniff evoked firing amplitude and an inhibition of firing rates outside the maximal sniff phase. These findings demonstrate that GABAergic neuromodulation affects MTC firing in a bimodal, sensory-input dependent way, suggesting that GABAergic basal forebrain modulation could be an important factor in attention mediated filtering of sensory information to the brain.

## Introduction

The basal forebrain (BF) is a complex of subcortical nuclei with projections to various brain areas and has been implicated in attention and cognitive control. It constitutes the primary source of cholinergic projections to limbic structures, the cortical mantle and olfactory areas ^1^. Cholinergic neuromodulatory systems are thought to enhance sensory processing and amplify the signal-to-noise ratio of relevant responses ^2-5^ e.g. the running-induced gain increases evident in sensory cortex ^6,7^ or the dishabituation of odor responses in the olfactory system ^8^. Furthermore, they have been identified as key players in mediating attentional modulation of sensory processing as well as in coordinating cognitive operations ^9,10^. However, the concept of a prevalent role of cholinergic cells in the BF was recently challenged as activity of non-cholinergic neurons was shown to strongly correlate with arousal and attention ^11-15^. Despite the knowledge of BF subpopulations containing neurotransmitters different from acetylcholine ^16-19^ no direct comparison of modulation effects caused by cholinergic and non-cholinergic projections is currently available. Especially for sensory processing which is strongly influenced by attentional states ^20^, effects of non-cholinergic BF modulation have been sparsely investigated.

The olfactory system in mice is heavily innervated by centrifugal inputs from the BF with the majority of bulbopetal neurons located in the horizontal limb of the diagonal band of Broca (HDB) ^21-26^. Though only about one fifth of BF neurons are cholinergic ^25^ studies on olfactory processing have mainly focused on cholinergic effects ^8,27-45^. In *in vivo* studies a specific activation of cholinergic HDB cell bodies was shown to inhibit spontaneous mitral tufted cell activity ^27^ while optogenetically activating cholinergic axons directly in the OB added an excitatory bias to OB output neurons: the enhancement of mitral/tufted cell odorant responses occurred independent of the strength or even polarity of the odorant-evoked response ^28^. The effect of cholinergic fiber stimulation is reminiscent of sensory gain modulation in the form of baseline control ^46^, which fits well to behavioral effects of nicotinic acetylcholine modulation in the OB ^36^ reported to increase behavioral discriminability.

Despite 30 % of the bulbopetal projections neurons in the BF being GAD-(glutamic acid decarboxylase, the rate-limiting enzyme in the synthesis of GABA) positive ^25^, less attention has been directed towards GABAergic BF OB projections ^23,47,48^. Using predominantly *in vitro* OB slice recordings, studies identified periglomerular interneurons ^48^ and granule cells ^47^ as targets of GABAergic projection.

Here, we used electrophysiological and optogenetic approaches to examine how cholinergic or GABAergic projections from BF modulate MTC output from the OB *in vivo*. We found marked differences between these projections; centrifugal cholinergic fibers from BF lead to an enhanced excitation of MTCs both at rest and in response to weak or strong sensory inputs. Effects of GABAergic BF axon stimulation in the OB on the other hand were sensory input strength dependent and mainly caused suppression of spontaneous MTC activity while predominantly enhancing odor evoked MTC spiking.

These results suggest that both, cholinergic and GABAergic projections from the same area, rapidly modulate sensory output but might have markedly different impacts on sensory information processing.

## Results

### Differential expression of ChR2 in basal forebrain projection neurons

To selectively target cholinergic or GABAergic projections from BF to the OB we used mouse lines expressing Cre under control of the ChAT (ChAT-Cre mice; ^49^) or the GAD2 promotor (GAD2-Cre, ^50^). We expressed channelrhodopsin specifically in cholinergic or GABAergic BF neurons using a Cre-dependent viral expression vector targeted to BF by stereotaxic injection (Supp. Fig. 1). As reported previously ^28,51,52^ viral injection in ChAT-Cre animals led to ChR2-EYFP expression on the somata and processes of neurons throughout HDB and, to a lesser extent, the vertical limb of the diagonal band of Broca (Fig. 1A). In few preparations sparsely labelled neurons could be additionally observed in the magnocellular preoptic nucleus (MCPO, data not shown). BF-injected GAD2-Cre animals displayed ChR2-EYFP expression predominantly in the HDB (Fig. 1B). In fewer cases the MCPO showed a sparser cellular expression.

**Figure 1.**
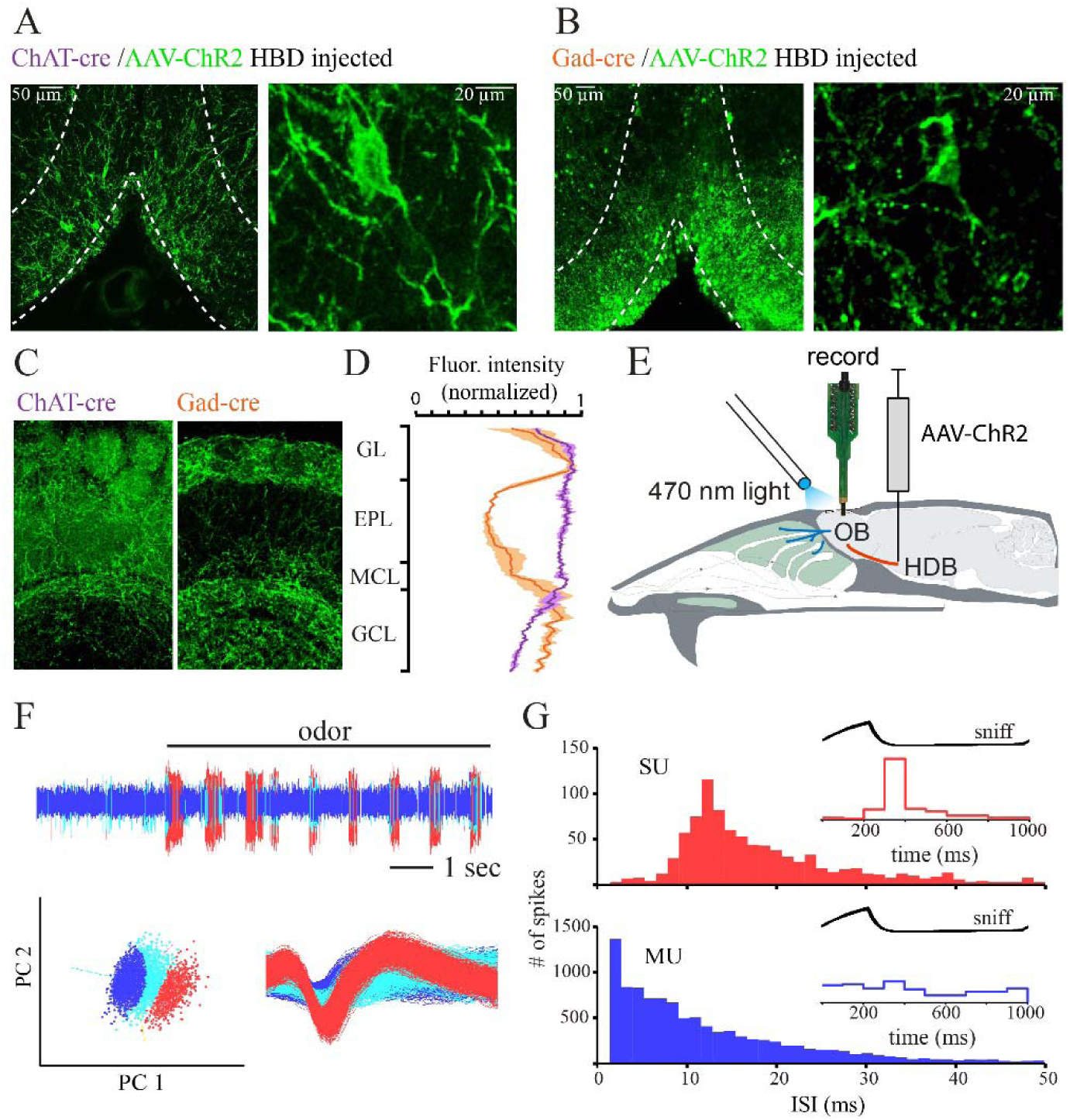
Selective targeting of cholinergic and GABAergic inputs from basal forebrain to the OB. A. Left, Coronal section (Bregma 0.74) through BF in a ChAT-Cre mouse injected with a Cre-dependent AAV-ChR2 virus. White lines indicate the outline of HDB and VDB. Right, Magnification of HDB showing labelled ChAT+ neurons. B. Left, Coronal section (Bregma 0.74) through BF in a GAD2-Cre mouse injected with a Cre-dependent AAV-ChR2 virus. White lines indicate the outline of HDB and VDB. Right, Magnification of HDB/VDB showing somata of GAD+ neurons. C. ChR2-EYFP-expressing axon terminals in different layers of the OB 4 weeks after AAV-ChR2-EYFP injection into BF of a ChAT-Cre and GAD-Cre mice. GL: glomerular layer, EPL: external plexiform layer, MCL: mitral and tufted cell layer, GCL: granule cell layer. D. Normalized fluorescence intensity profiles show that axons reaching the external plexiform layer are less prominent in GAD-Cre mice. E. Schematic of experimental approach. See Materials and Methods for details. F. Data acquisition. In continuous recordings action potential waveforms with a signal-to-noise ratio of at least 4 SD above baseline noise were saved to a disk (sample rate 24 kHz) and further isolated using off-line spike sorting. Vertical line indicates onset of odorant stimulation. Isolation of waveforms into three different units by principle components 1 and 2 (bottom, left). Spike waveforms of the isolated units (bottom, right). G. Inter-spike-interval histograms for two units. One is a single unit (SU) and one a multi unit (MU). Only single units were analysed further. For units to be classified as presumptive MTCs also a clear sniff-modulation in sniff triggered spike averages had to be present (inset).

Four weeks after virus infection, ChR2-EYFP protein was apparent in BF fibers throughout the OB (Fig. 1C, D). In ChAT-Cre mice labelled axon terminals were visible in all layers of the OB (Fig. 1C left, Fig. 1D), consistent with earlier reports about cholinergic fibre distribution ^25,28,53-56^. The fluorescence intensity of EYFP per area unit was uniform across higher OB layers and declined in the granule cell layer. In GAD2-Cre animals the OB was also densely innervated by labelled fibres. Here, the fluorescence intensity per area unit was especially high in the glomerular and the granule cell layer (Fig. 1C right, Fig 1D) recapitulating previous finding ^47^ but also strong in the mitral cell layer. Fluorescence intensities were distinctly lower in the external plexiform layer, the main location of MTC / GC dendrodendritic synapses. The normalized fluorescence intensity per area unit of single fibers in ChAT-Cre and GAD-Cre OB was not significantly different (1.00 ± 0.06 and 1.06 ± 0.06, respectively; n = 3 mice, p = 0.52). Therefore, the differences in average fluorescence intensities reflect the difference in fiber density rather than ChR2-EYFP expression levels.

### Optogenetic activation of cholinergic and GABAergic axons in the OB modulates MTC spontaneous spiking

To investigate BF modulation effects on early olfactory processing, we directed 473 nm light (1-10 mW total power) onto the dorsal OB surface while recording multi-channel electrical activity from dorsally located presumptive MTCs in anesthetized, double-tracheotomized mice (Fig. 1E-G) (see Materials and Methods).

To access the impact of cholinergic and GABAergic fiber stimulation on MTC excitability in the absence of sensory input, we optically activated BF axons without ongoing inhalation (Fig. 2A). In this condition, MTC display an irregular firing pattern ^28,57,58^. As shown previously ^28^, optogenetic activation of cholinergic fibers in ChAT-Cre mice lead to a significant increase of spontaneous MTC spiking from 2.05± 2.34 Hz (mean ± SD) before stimulation to 2.40± 2.24 Hz during stimulation (n = 27 units from 5 mice; *p* = 0.0157 Wilcoxon signed rank test). 8 of these units (30%) showed a significant stimulation-evoked increase in firing activity when tested on a unit-by-unit basis (Mann–Whitney *U* test); none showed a decrease (Fig. 2B, left).

**Figure 2:**
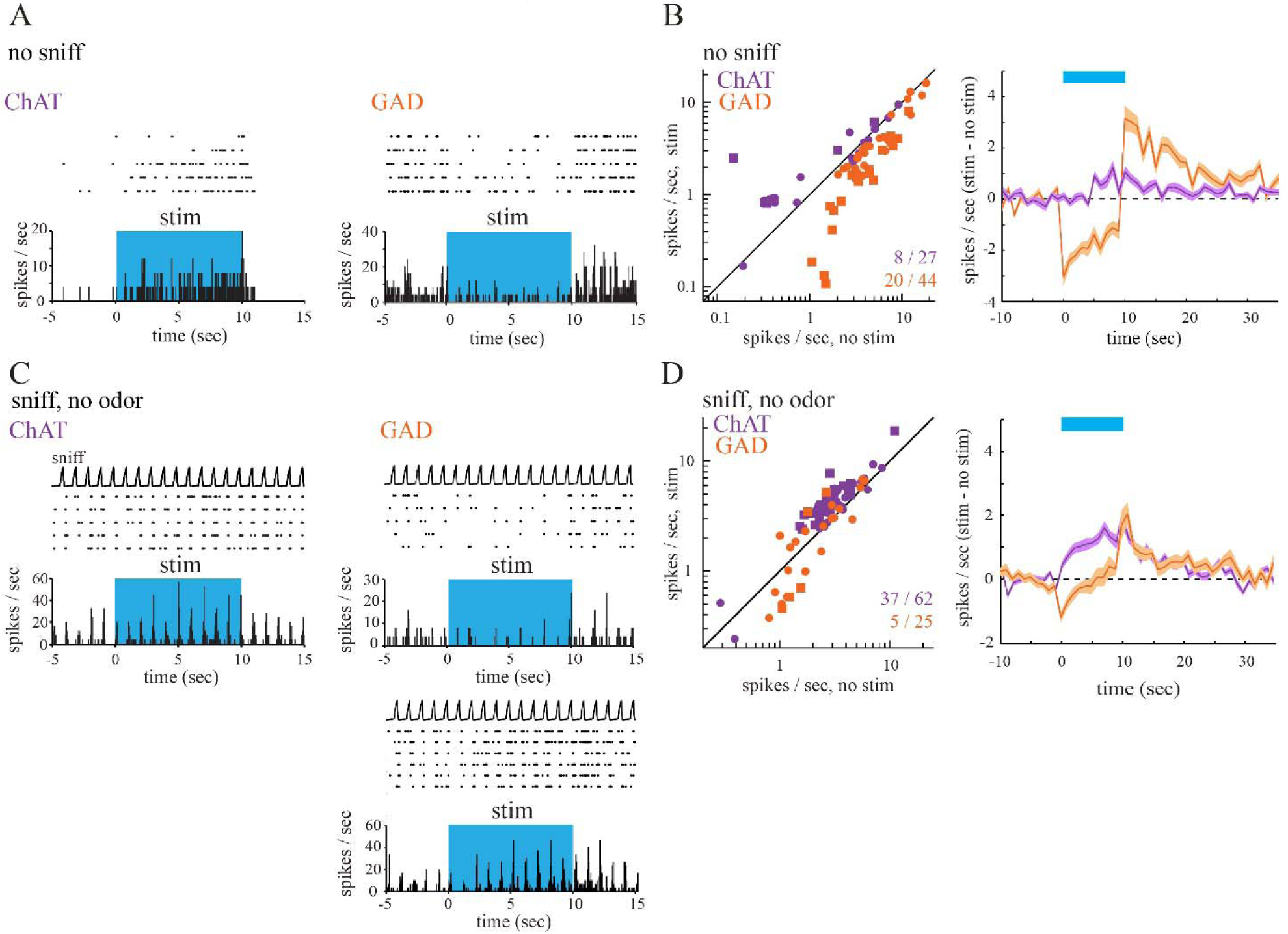
Optogenetic activation of cholinergic and GABAergic basal forebrain inputs to the OB modulates spontaneous as well as sensory-evoked MTC spiking. A. Spike rasters and rate histograms (bin width, 50 ms) from presumptive MTCs showing spontaneous spiking in the absence of inhalation (no sniff). Spike rate decreased during optical stimulation of the dorsal OB (“stim”, blue shaded area) in GAD-Cre mice and increased in ChAT-Cre animals. B. Left, Plot of spontaneous firing rate in the 9 s before (no stim) and during (stim) optical stimulation for all tested units (ChAT-Cre, n= 27 units, purple; GAD-Cre, n= 44 units, orange). Squares indicated significantly modulated units subjected to a unit-by-unit test. Right, Time course of change in firing rate (mean ± SEM across all units) during optical stimulation (blue bar). The trace indicates change in mean spike rate in 1 s bins relative to the mean rate before stimulation. The time axis is relative to time of stimulation onset. C. Spike raster and rate histogram of MTC spiking during inhalation of clean air and optical stimulation (blue shaded area). Inhalation-evoked spike rates decreases (top) or increase (bottom) during optical stimulation in GAD-Cre mice while only excitation in response to optical stimulation is observed in ChAT-Cre mice. The top trace (sniff) shows artificial inhalation as measured by a pressure sensor connected to the nasopharyngeal cannula. D. Left, Plot of inhalation-evoked firing rates during clean air inhalation, averaged for the nine inhalations just before (no stim) and after (stim) optical stimulation (ChAT-Cre, n= 62 units, purple; GAD-Cre, n= 25 units, orange). Data were analyzed and plotted as in B. Squares indicate significantly modulated units. Right: Time course of change in firing rate (mean ± SEM across all units) during optical stimulation (blue bar).

In contrast, optical stimulation in GAD2-Cre mice led to a significant decrease in MTC spontaneous spiking, from 5.43 ± 4.07 Hz (mean ± SD) before stimulation to 3.45 ± 3.54 Hz during stimulation (n =44 units from 5 mice; *p* = 1.07 × 10^−8^, Wilcoxon signed rank test, Fig. 2A, B). When tested on a unit-by-unit basis 20 of the 44 recorded units showed a significant reduction in firing activity while none showed a significant increase (Fig. 2B, left). The median reduction in spike rate across these cells was 1.78 ± 1.36 Hz. Across the population of all recorded units, the decrease in spontaneous firing rate persisted for the duration of the 10 s optical stimulation (Fig. 2B, right). Following the stimulation, an increase in spiking was observed that returned to prestimulation levels within 20 s after stimulation ceased. Thus, optogenetic activation of GABAergic BF fibers at the level of the OB leads to a reduction of MTC activity while activating cholinergic fibers causes output neuron excitation, demonstrating that these subpopulations cause opposing effects on spontaneous MTC firing.

In order to rule out optical activation artifacts we stimulated the OB of uninjected control mice with the same parameters as before during the no sniff condition, since in this condition even small changes could have been detected (Supp. Fig. 2). We found that optical stimulation led to no significant change in spontaneous firing rate (n = 19 units from three mice; 6.10 +-4.15 Hz before stimulation, 6.20 +-4.25 Hz during stimulation (mean ± SD); *p* = 0.365, Wilcoxon signed rank test). Thus, the light-evoked modulation of spontaneous MTC firing in ChR2-injected ChAT-Cre and GAD-Cre mice was attributable to cholinergic/GABAergic signaling mediated by the optogenetic activation of cholinergic/GABAergic axons in the OB.

### Optogenetic activation of cholinergic and GABAergic axons in the OB modulate inhalation-evoked MTC spiking

Next, we investigated the effect of cholinergic and GABAergic axon activation on MTC responses during artificial inhalation of clean air (Fig. 2C, D). Inhalation-linked spiking pattern could be observed in 62 units (4 mice) in ChAT-Cre and 25 units (6 mice) in GAD-Cre mice, most likely reflecting weak sensory-evoked responses ^28,59,60^.

Optical stimulation of cholinergic axons in ChAT-Cre mice significantly increased inhalation-linked spiking of MTCs (Fig. 2C), with median spike rate increasing from 2.86 ± 1.72 Hz to 4.03 ± 2.45 Hz during optical stimulation (p = 2.99 × 10^−11^, Wilcoxon signed rank test). When tested on a unit-by-unit basis, 37 of 62 recorded units (60%) showed significant optical stimulation evoked increase in spiking (Fig. 2D); none showed a decrease. Inhalation evoked spiking of these 37 cells increased by 1.33 ± 1.29 spikes/ sniff/s.

In GAD2-Cre mice optical stimulation led to mixed effects on inhalation-linked spiking that were not significant across the population of MTCs (1.8 ± 1.66 Hz before stimulation, 2.26 ± 2.13 Hz during stimulation; n = 25 units from 6 mice; *p* = 0.58 Wilcoxon signed rank test, Fig. 2C, D). When tested on a unit-by-unit basis three of the 25 (12%) units showed a significant decrease and two (8%) a significant increase in firing activity. Inhalation evoked spiking decreased by 0.67 ± 0.14 spikes/ sniff/s and increased by 2.35 ± 0.71 spikes/ sniff/s for the significantly inhibited and excited units, respectively. The averaged time course depicted an initial decrease in MTC firing rate that, in contrast to the time course in the no sniff condition, returned to prestimulation levels already during the stimulation period (Fig. 2D). Following stimulation, spike rate increased above baseline levels for approx. 20 s before returning to baseline. Taken together, while optogenetic stimulation of cholinergic fibers in both conditions was qualitatively similar, activating GABAergic fibers leads to mixed effects of MTC spiking in the sniff condition that were not observed during spontaneous spiking.

### Optogenetic activation of cholinergic and GABAergic axons in the OB enhances odorant-evoked MTC spiking

Since optical activation of BF inputs to the bulb modulates inhalation-linked MTC spiking consistent with modulating weak sensory-evoked responses, we next evaluated the impact of bulbar cholinergic and GABAergic modulation on odorant responses. We compared MTC responses to odorant stimulation applied with and without optogenetic activation of cholinergic or GABAergic BF axons (Fig. 3A, B). In line with the findings from the previously tested conditions light activation of cholinergic fibers in ChAT-Cre mice increased MTC spiking (Fig. 3A, C). Across the population of recorded presumptive MTCs (n= 56 units, 5 mice), BF axon stimulation significantly increased MTC odor activity, with an increase from 6.49 ± 3.47 spikes/sniff/s (median ± SD) during odorant presentation alone to 8.98 ± 4.22 spikes/sniff/s during odorant paired with light (*p*=3.93 × 10^−10^, Wilcoxon signed rank test). 25 out of the 56 recorded cells (45%) showed a significant increase in odor-evoked spiking when tested on a unit-by-unit basis; none showed a decrease (Fig. 3C).

**Figure 3:**
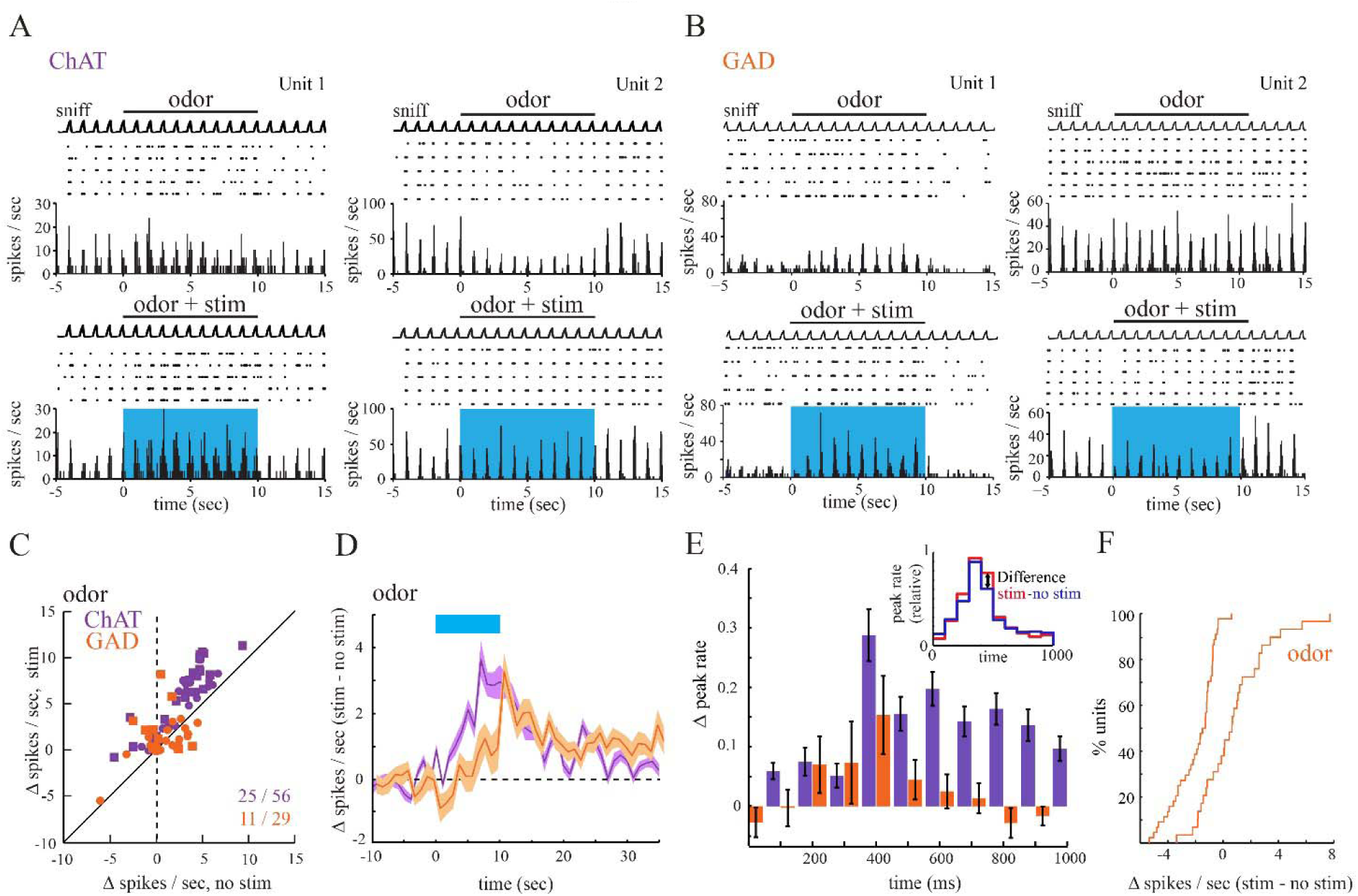
Optogenetic activation of cholinergic and GABAergic OB inputs modulates odor-evoked responses. A. Odorant-evoked MTC spiking is enhanced by optical OB stimulation in ChAT-Cre mice in both odorant-activated (Unit1) and odorant-inhibited (Unit2) cells. B. Odorant-evoked MTC spiking is increased (left) or decreased (right) by optical OB stimulation in GAD-Cre mice. C. Plot of odorant-evoked changes in MTC spiking (Δ spikes/sniff) in the absence of (no stim) and during (stim) optogenetic stimulation of cholinergic (n = 56 units, purple) or GABAerig (n = 29, orange) afferents to the OB. D. Time course of effects of optical stimulation on odorant-evoked spike rate, averaged across all units. The blue bar shows time of optical stimulation and simultaneous odorant presentation. Plotted is the mean change in odorant-evoked spike rate between trials with and without light stimulation, measured after each inhalation (at 1 Hz); the shaded area indicates variance (SEM) around mean. E. Optical stimulation induced firing changes during the sniff cycle. Cholinergic modulation increased the firing rate across the sniff cycle. GABAergic modulation increased firing in peak and adjacent time bins while inhibiting firing outside the “preferred sniff phase”. Firing changes were calculated form sniff triggered spike histograms as depicted in the inset. F. Cumulative probability plots comparing change in MTC firing rates caused by optical stimulation in the no sniff and odor condition. Plots reflect datasets plotted in Fig. 1B and Fig. 3C, respectively. Unlike the uniform suppression of spontaneous spiking observed in GAD-Cre mice, optical stimulation in the odor condition predominantly causes MTC excitation.

Optogenetic activation of GABAergic fibers in the OB during odor presentation elicited heterogeneous effects that, across the population of recorded cells (n= 29 units, 4 mice), were not significant (5.44 ± 3.12 spikes/sniff/s during odorant presentation alone, 4.93 ± 4.30 spikes/sniff/s during odorant paired with light, *p* = 0.52, Wilcoxon signed rank test). However, tested on a unit-by-unit basis eight out of 29 recorded cells (28 %) showed a significant increase and three (10 %) a significant decrease in odor-evoked firing activity (Fig. 3C). The modulation in spike rate across the cells showing a significant increase /decrease was 2.17 / 2.48 spikes/sniff/s (from 7.53 ± 4.1 to 9.7 ± 5.32 / from 5.44 ± 0.57 to 2.96 ± 0.39), respectively. The averaged activity time course of the recorded units displayed an initial brief reduction of firing rated that switched to excitation during the stimulation period (Fig. 3D).

We also examined optical stimulation induced changes in odor-evoked MTC spiking pattern by generating sniff triggered spike histograms aligned to the start of inhalation for ChAT-cre and GAD-cre mice (Fig 3E inset, see ^28^. Neither cholinergic nor GABAergic basal forebrain derived modulation showed a significant change of the time bin of peak firing across the population of recorded units (*p* = 0.63 and 1 for ChAT-cre and GAD-cre mice, respectively, paired *t* test). However, comparing the optical stimulation induced firing change per bin revealed a profound difference between cholinergic and GABAergic modulation (Fig. 3E): while cholinergic modulation increased the firing rate across the sniff cycle, GABAergic modulation increased firing in peak and adjacent time bins while inhibiting firing outside the “preferred sniff phase”.

The BF receives input from different olfactory areas ^45^ and it has been shown that even during sleep and anesthesia, cholingergic and GABAergic BF neurons are rhythmically discharging ^61-64^. We therefore tested the effect of inhibiting cholinergic and GABAergic BF projections to the OB using the light-gated chloride pump Halorhodopsin as an optogenetic silencer (Supp. Fig. 3). Despite robust, yet sparser expression of Halo-YFP, labelling could be observed in the OB for both ChAT-Cre and GAD-Cre mice four weeks after viral injection (Supp. Fig. 3A). Optogenetic stimulation during recording of presumptive MTCs (Supp. Fig. 3B) showed, when tested on a unit-by-unit basis, no significant effects in ChAT-Cre animals (8 units, 2 mice) while only two out of 28 recorded cells showed a weak but significant decrease in odor-evoked spiking (0.47 spikes/sniff/s) in GAD-Cre animals (Supp. Fig. 3C). No significant modulation effects were observed in the other tested conditions (no sniff and sniff, data not shown). This was also true for units showing strong sensory evoked spiking, rendering it unlikely that effects went undetected due to low spike counts. The surprisingly weak effects of optogenetic inhibition might be the result of only weak spontaneous cholinergic and GABAergic OB fibre activity in the anesthetized animal.

In uninjected control mice (Supp. Fig. 2), the same optical stimulation led to no significant change in spontaneous firing rate (n = 11 units from three mice; 6.31 +-5.56 Hz before stimulation, 6.26 +-5.47 Hz during stimulation (mean ± SD); *p* = 0.61, Wilcoxon signed rank test).

### GABAergic projections modulate OB output dependent on sensory input

Unlike spontaneous spiking, which got suppressed by optical stimulation in GAD-Cre mice, the same optical stimulation during odor stimulation predominantly caused MTC excitation (Fig. 3 F). In a separate set of experiments we therefore investigated if this suppression to excitation transition can also be observed on a single unit basis, or might be caused by a recording bias e.g. through different populations of output neurons being detectable in the different conditions. We recorded GABAergic modulation effects in individual MTC tested in both the spontaneous as well as the odorant evoked condition in one continuous session (Fig. 4 A and B depict recordings from the same unit).

**Figure 4.**
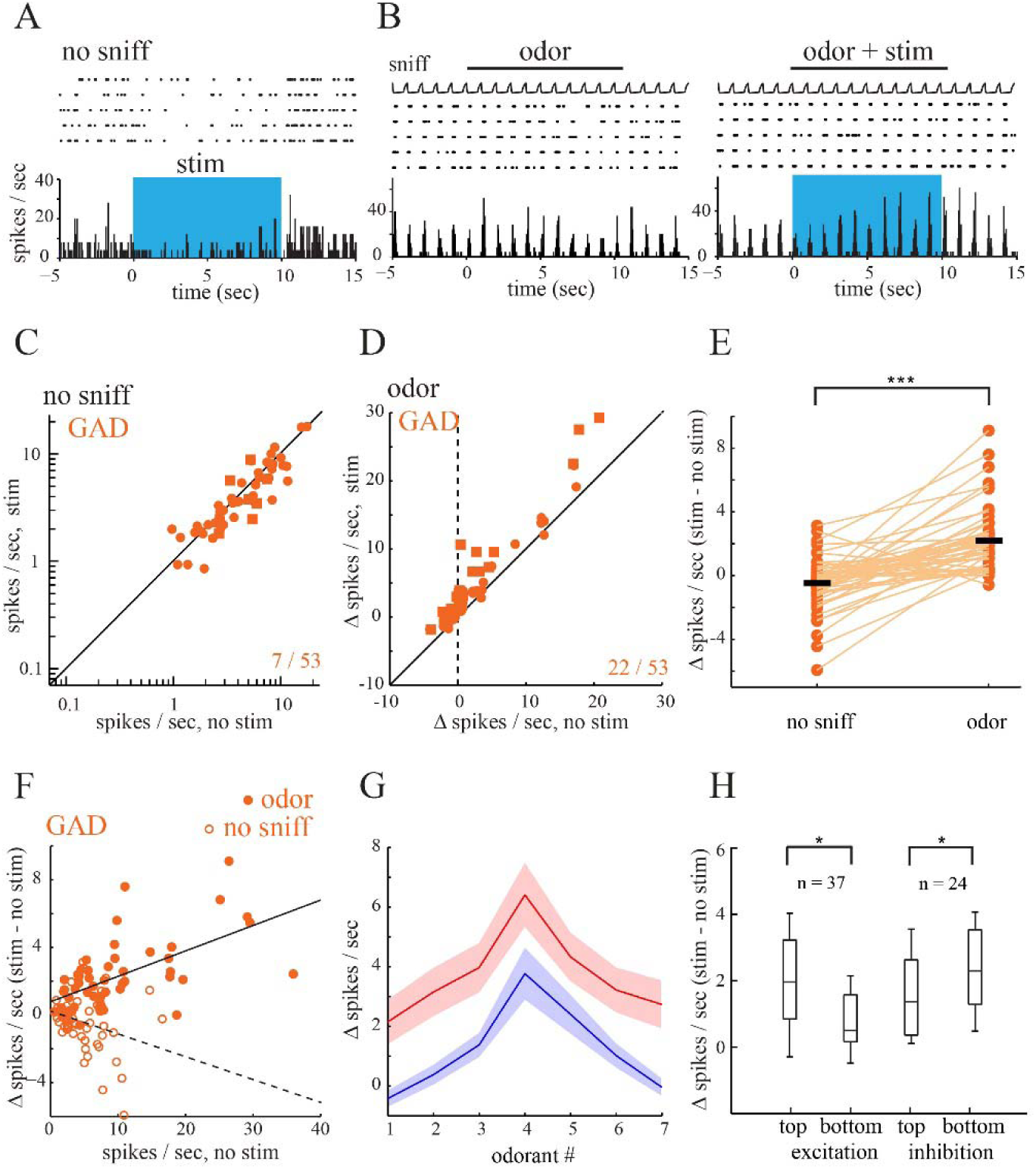
Switch in modulatory effect between conditions in GAD-Cre mice. A. Spike raster and rate histogram (bin width, 50 ms) depicting a spike rate decrease for a MTC during optical stimulation of the dorsal OB (“stim”, blue shaded area). B. The same unit plotted in A was also tested in the odor condition. Odorant-evoked spiking is enhanced by optical OB stimulation in this unit. C. Plot of spontaneous firing rate in the 9 s before (no stim) and during (stim) optical stimulation for all units tested in both the no-sniff and the odor condition (n= 53 units). Squares indicated significantly modulated units subjected to a unit-by-unit test. D. Plot of odorant-evoked spiking changes (Δ spikes/sniff) in the absence of (no stim) and during (stim) optogenetic stimulation of the same units tested in C. E. Quantitative comparison of stimulation-evoked spiking changes (Δ spikes/sec) in the no-sniff and odor condition. Open circles, spiking changes for individual units; filled bars, mean value. Lines connect the same unit across conditions. F. Optical stimulation effects in the odor condition were positively correlated to baseline activity (Pearson’s *r* = 0.61; two-tailed, *p* = 1.47 × 10^−6^). A negative correlation was observed in the no-sniff condition (Pearson’s *r* = −0.27; two-tailed, *p* = 0.05). G. Effect of optical GABAergic OB fiber stimulation on odorant response spectrum for MTCs tested with seven (right) odorants. Blue, baseline response; red, response during optical stimulation. Odorants are ordered separately for each unit, with the strongest excitatory response in the baseline condition in the middle of the abcissa. Lines (±SEM) connect median responses across all tested trials. H. Box plot comparing optically induced change in odorant-evoked spike rate (stim - no stim) for the strongest and weakest quartile of baseline odorant-evoked responses taken from all cell– odorant pairs for excitatory as well as inhibitory responses. The box indicates median, 25–75th percentile ranges of the data, and whiskers indicate ±1 SD from the mean.

As shown previously (Fig. 2), optical stimulation in the no sniff condition led to a significant reduction in MTC spontaneous spiking across all recorded units (3.61 ± 3.43 Hz before stimulation; 3.14 ± 3.37 Hz during stimulation; n =53 units from 3 mice; *p* = 0.014 Wilcoxon signed rank test). When tested on a unit-by-unit basis 5 of the 53 units recorded showed a significant reduction in firing activity and only two units showed an increase (Fig. 4C). Similar to the previous findings (Fig. 3), GABAergic axon activation during odor presentation had a predominantly excitatory effect on MTC activity (7.35 ± 8.12 spikes/sniff/s during odorant presentation alone, 8.13 ± 9.47 spikes/sniff/s during odorant paired with light, *p* = 9.86 × 10^−10^, Wilcoxon signed rank test). Moreover, all significantly modulated units (22 out of 53 cells (42%)) showed an optical stimulation induced increase in odor-evoked spiking, none showed a decrease. Quantitative comparison of the optogenetic stimulation effect in both conditions showed a significantly more positive modulation in the odor compared to the spontaneous condition (Fig. 4E; spontaneous spiking −0.31 ± 1.61 vs. odor-evoked spiking 1.98 ± 2.01 (median ± SD); measured as Δ spikes/s relative to non-stimulation condition; *p* = 6.34 × 10^−9^, Wilcoxon signed rank test). Comparing the firing rate of individual units between the conditions revealed that almost all recorded units (48 of 53 units) showed a more positive, optically evoked modulation in spike rate in the odor compared to the no sniff condition.

In order to investigate in how far the observed modulation effects depend on baseline firing rates we plotted the optical stimulation effect as a function of baseline activity. As expected, baseline firing activity was lower in the no-sniff compared to the odor condition (3.61 ± 3.43 (median ± SD) spikes/sec compared to 7.35 ± 8.11 spikes/sec; *p* = 2.53 × 10^−6^, Wilcoxon signed rank test). Interestingly, while optical stimulation effects in the odor condition were positively correlated to baseline activity (Pearson’s *r* = 0.61; two-tailed, *p* = 1.47 × 10^−6^) a negative correlation was observed in the no-sniff condition (Pearson’s *r* = −0.27; two-tailed, *p* = 0.05). These results suggest that stimulation effect most likely depend on the amount of sensory input rather than solely on the absolute firing rate prior to optical stimulation. The results furthermore indicate that the inhibition to excitation transition across conditions can also be observed on a single unit basis and thus that GABAergic fiber derived modulation can act in a bimodal way on single OB output neurons.

Previous studies have suggested a sharpening of MTC odorant responses by cholinergic modulation through preferentially suppressing weak MTC responses and enhancing inhibitory responses ^27,33,43^. Since we only observed additive cholinergic modulation effects on MTC activity, suggesting no change of tuning in accordance to a previous report ^28^, we asked if the GABAergic basal forebrain subsystem might be capable of sharpening odor responses in the OB. In a separate set of experiments (36 units, from 3 animals) we therefore tested the response of the same MTC to multiple odorants with and without optical GABAergic OB fiber stimulation. Consistent to our previous findings, we found that odorant responses were mostly enhanced (0.39 ± 3.22 spikes/sniff/s during odorant presentation alone, 2.41 ± 5.01 spikes/sniff/s during odorant paired with light, *p* = 1.79 × 10^−40^, Wilcoxon signed rank test). Stimulation effects on tuning curves varied between units. An average tuning curve, calculated across the population of all recorded units (Fig. 4G) displayed a uniform, odor strength independent increase in firing rather than a change in tuning. However, when odorant– cell responses were collapsed across units creating a large dataset of 252 cell odor pairs (Fig. 4H), a comparison of the optically induced change in odorant-evoked spike rate (stim-no stim) for the strongest and weakest quartile of odorant-evoked responses revealed significant differences for excitatory (unpaired Wilcoxon rank-sum test, p = 0.03; n = 37 cell–odor pairs) as well as inhibitory (unpaired Wilcoxon rank-sum test, p = 0.01; n = 24 cell–odor pairs) responses. These results points to strong odorant responses receiving more excitatory modulation from GABAergic basal forebrain derived fibers no matter if their particular odorant response is excitatory or inhibitory.

## Discussion

The basal forebrain is critical for many cognitive processes ^65-69^ as well as for sensory information processing ^51,70-75^. Despite longstanding knowledge about the heterogeneity of BF derived long range projections ^17,25,76,77^, studies examining their function in sensory processing so far have been focused almost exclusively on cholinergic projections ^3,27,28,46,78-82^.

Here, we examine the effect of GABAergic long range projections in the olfactory bulb (OB) and provide for the first time a direct comparison of cholinergic and GABAergic modulation effects on early sensory processing under the same experimental conditions.

### Expression and activation of channelrhodopsin in basal forebrain projections

Differential expression of channelrhodopsin in either GABAergic or cholinergic long range projections was achieved by injection of AAV-ChR2 into the BF of GAD2 or ChAT-cre mice, respectively. Expression patterns showed a relatively homogenous distribution of cholinergic fibers in the OB in accordance with previous publications. Distribution of GABAergic fibers only differed slightly from a previous publication that used the same transgenic mouse line with a similar expression strategy ^47^. Similar to their results we saw dense labeling in the granule cell layer, but equally strongly labelled glomerular layer and a slightly weaker labelled mitral cell layer. This slight difference could be the result of viral injection sites since Nunez-Parra et al. targeted more posterior parts of BF.

A recent report points to a potential coexpression of cholinergic and GABAergic markers at the level of the OB ^83^. Since transgene expression was obtained by crossing Cre driver lines to reporter animals, coexpression might have occurred at some point during development, not reflecting the adult situation. The different expression patterns and photostimulation effects of cholinergic and GABAergic BF derived projections observed here, suggest that these projections are mainly separated and ACh and GABA are not likely being released from the same fibers. In line with our findings, a report using the same GAD2-Cre line as used in the present study, did not observe any non-GABAergic currents upon stimulation of BF derived GABAergic fibers ^47^.

Activation of long range projections was performed by photostimulating axonal fibers at the dorsal surface of the OB. While known to be less effective compared to somatic stimulation ^84^, optogenetic fiber stimulation at the level of the OB was necessary to obviate potential indirect BF modulation effects of OB activity: First, optogenetically activating BF might cause a stimulation of brain areas also targeted by BF fibers that in turn project to the OB ^25,76,77,85-91^. Second, a direct BF stimulation in GAD-Cre mice would cause activation of the large amount of GAD positive inhibitory BF interneurons ^92,93^. Third, cholinergic BF neurons were shown to excite GABAergic projection neurons ^94-96^, thereby also rendering a direct BF stimulation problematic. Axonal fibers at the dorsal surface of the OB were activated using a continuous stimulation paradigm, which has been shown to be most effective in activating cholinergic axonal fibers ^28,97^. BF fiber silencing with inhibitory opsins hardly showed any MTC activity modulation in the different conditions (sniff, no sniff, odor). The surprisingly weak effects of optogenetic inhibition might be the result of only weak spontaneous cholinergic and GABAergic OB fibre activity in the anesthetized animal.

### Effects of basal forebrain projections on OB output cell activity

Recording from olfactory bulb output neurons, we show that, in contrast to a local and specific activation of cholinergic fibres, that add an excitatory bias to mitral/tufted cell firing, a selective activation of GABAergic BF fibres leads to bimodal, sensory input dependent effect on OB output: whereas optogenetic stimulation mainly inhibited spontaneous MTC firing, odor evoked MTC cell spiking was predominantly enhanced; an effect that could also be observed on a single neuron level. Additionally, MTCs showed a reduction of firing outside and an increase of firing within the preferred sniff phase. This modulation is strongly reminiscent of a model of a bimodal gain change evoked by attention ^9^ also referred to as filtering. These filter processes are reported to dampen activity to non-attended or background stimuli while enhancing relevant sensory input. Indeed, in the odor condition, the size of the GABAergic modulation effect was dependent on the odor evoked firing activity for both excitatory and inhibitory odor responses, pointing to a multiplicative population firing change for relevant olfactory stimuli while decreasing background activity.

Our direct comparison of cholinergic and GABAergic OB fiber activation suggest that both BF derived fiber systems might have a role in gain modulation of OB output, but in a very distinct way: while cholinergic modulation seems to be rather similar to baseline control ^46^, GABAergic modulation seems to lead to a filtering of weak signals in the OB.

### Possible circuit mechanisms underlying bulbar GABAergic modulation

The circuit mechanisms underlying basal forebrain derived modulation effects remain to be elucidated since a comprehensive list of targets for BF fibers in the OB is missing especially for GABAergic projections. Cholinergic and GABAergic systems most likely exert multiple effects within the OB with different receptors, cell types and modes of transmission (volume vs. strict synaptic transmission) involved ^98^.

Just like in several other brain areas ^99^, GABAergic BF axons in the OB are so far only reported to synapse on inhibitory interneurons ^23,47^: centrifugal GABAergic afferents were shown to inhibit granule cells ^47^ deep short axon cells ^48,83^ as well as different types of periglomerular cells ^48^. Inhibition of those inhibitory interneuron subtypes would, in the simplest case, result in a disinhibition of mitral and tufted cells and could therefore explain the here observed MTC excitation. The sensory input dependent increase in excitatory modulation between conditions and the correlation of effects size with odorant evoked responses could be the result of an increase in sensory dependent activity in OB interneurons ^100^ leading to a further increase in disinhibition (Suppl. Fig. 4). The observed inhibition in MTCs could be elicited by a multisynaptic effect via several types of inhibitory interneurons: neurons like deep short axon cells and PG cells (which receive GABAergic BF input ^48^) were shown to inhibit other inhibitory interneurons like periglomerular and granule cells, respectively (see ^101^). However, the fast, strong and persistent nature of this inhibition, rather argues for a direct inhibitory input to MTCs Suppl Fig. 4) similar to that observed in the basolateral amygdala where BF GABAergic fibers provide indirect disinhibition, as well as direct inhibition of pyramidal neurons ^102^.

## Conclusion

Our recordings provide, for the first time, a detailed comparison of BF cholinergic and GABAergic influence on early sensory processing and highlight the potential of the noncholinergic BF population to modulate perception. By inhibiting weak and facilitating strong inputs, GABAergic BF fibres in the OB likely increase MTCs signal-to-noise ratio, a hallmark of attentional processes that have been previously attributed mainly to cholinergic processes ^2-4,9,10^. Our findings are in line with recent data indicating that the classical view on the (cholinergic) BF system might be oversimplified: activity of non-cholinergic BF neurons was more strongly correlated with arousal and attention ^11-15^, whereas cholinergic neuron activity was correlated with body movements, pupil dilations, licking, punishment ^75,103^ as well as primary reinforcers and outcome expectations ^11^. The distinct early sensory modulation effects of cholinergic and GABAergic BF neurons observed in this study might therefore be owed to the different functions of these two basal forebrain systems. Addressing the exact interplay between ACh and GABA in olfactory bulb sensory processing in the awake animal will be critical to fully understand the relative contribution of each system.

## Materials and Methods

### Animals strain and care

We used mice expressing Cre recombinase under control of the choline acetyltransferase (ChAT-Cre mice; JAX Stock #006410, The Jackson Laboratory) or glutamate decarboxylase 2 (GAD2-Cre mice, JAX Stock #010802, The Jackson Laboratory) promotor ^49,50^. Animals of either sex were used. Animals were housed under standard conditions in ventilated racks. Mouse colonies were bred and maintained at RWTH Aachen University animal care facilities. Food and water were available *ad libitum*. Experimental protocols were approved by the “*Landesamt für Natur, Umwelt und Verbraucherschutz NRW (LANUV NRW) (State Office for Nature, Environment and Consumer Protection North Rhine-Westphalia), Postfach 10 10 52, 45610 Recklinghausen, Germany”* and are in compliance with European Union legislation and recommendations by the Federation of European Laboratory Animal Science.

### Viral vectors

Viral vectors were obtained from the viral vector core of the University of Pennsylvania or Addgene. Vectors were from stock batches available for general distribution. pAAV-EF1a-double floxed-hChR2(H134R)-EYFP-WPRE-HGHpA and pAAV-Ef1a-DIO eNpHR 3.0-EYFP were a gift from Karl Deisseroth (Addgene viral prep # 20298-AAV1; http://n2t.net/addgene:20298; RRID: Addgene_20298 and Addgene viral prep # 26966-AAV1; http://n2t.net/addgene:26966; RRID:Addgene_26966)). Injection of Cre-dependent vector (AAV1.EF1a.DIO.hChR2(H134R)-eYFP.WPRE.hGH (*AAV.FLEX.ChR2.YFP*) and AAV1.EF1a.DIO.eNpHR3.0-eYFP.WPRE.hGH (*AAV.FLEX.HR.YFP*)) was performed as described in ^28,104^. Briefly, BF virus injection in adult (≥ 8 weeks) homozygous ChAT-Cre or GAD2-Cre mice was performed using stereotaxic targeting (relative to Bregma (in mm) +0.74 anteroposterior, 0.65 mediolateral, −4.8 dorsoventral, ^105^). Virus (0.5 - 0.75 μl; titer 1.97 × 10^12^ - 2.35 × 10^12^) was delivered through a 26 gauge metal needle at a rate of 0.1 μl/min. Mice were individually housed for at least 28 days before evaluating for transgene expression or recording.

### Olfactometry

Odorants were presented as dilutions from saturated vapor in cleaned, humidified air using a custom olfactometer under computer control ^28,106,107^. Odorants were typically presented for 10 seconds. All odorants were obtained at 95 - 99% purity from Sigma-Alrich and stored under nitrogen/argon. The following odorants were used: ethyl butyrate, 2-hexanone, methyl valerate, valeraldehyde, methyl hexanoate, isoamyl acetate, sec-butyl acetate, vinyl butyrate and ethyl tiglate. Odorants were presented at 1% saturated vapor (s.v.).

### Extracellular recordings and optical stimulation

MTC unit recordings and optical OB stimulation were performed as described previously ^28^ with several modifications. Briefly, mice were anesthetized with pentobarbital (50 mg/kg) and placed in a stereotaxic device. Mice were double tracheotomized and an artificial inhalation paradigm used to control air and odorant inhalation independent of respiration ^108-110^. Extracellular recordings were obtained from OB units using sixteen channel electrodes (NeuroNexus, A1×16-5mm50-413-A16, Atlas Neuro, E16+R-100-S1-L6 NT) and an RZ5 digital acquisition system (TDT, Tucker Davis Technologies). Recording sites were confined to the dorsal OB. Action potential waveforms with a signal-to-noise ratio of at least 4 SD above baseline noise were saved to a disk and further isolated using off-line spike sorting (Open-Sorter; TDT, Fig. 1F). Sorting was done using the Bayesian or (in fewer cases) K-Means cluster cutting algorithms in OpenSorter. Units were defined as “single units” if they fell within discrete clusters in a space made up of principle components 1 and 2. Units with interspike intervals lower than the absolute refractory period (< 2.5 ms) were excluded from further analysis ^111^ (Fig. 1G). For units to be classified as presumptive MTCs, units additionally had to be located in the vicinity of the mitral cell layer, show spiking activity in the absence of odorants, and a clear sniff-modulation (the maximum spike rate in a 100 ms bin had to be at least two times the minimum spike rate for the PSTH (Fig. 1G insets)); similar as described in ^57^. Subsequent analyses were performed using custom scripts in Matlab. Odorant alone (‘baseline’) and odorant plus optical stimulation trials (at least 3 trials each) were interleaved for all odorants (inter-stimulus interval 40-50 s). Recordings were subject to unit-by-unit statistical analysis as described below.

For optical OB stimulation, light was presented as a single 10 - sec pulse either alone or simultaneous with odorant presentation using a 470 or 565 nm LED and controller (LEDD1B, Thorlabs) and a 1 mm optical fiber positioned within 3 mm of the dorsal OB surface as in earlier studies ^28^. The light power at the tip of the fiber was maximal 3 and 10 mW for the 565 nm and 470 nm LED, respectively.

### Extracellular Data analysis

Basic processing and analysis of extracellular data followed protocols previously described for multichannel MTC recordings ^28^. Responses to optical or odorant stimulation were analyzed differently depending on the experimental paradigm. Stimulation effects on spontaneous spike rate (no artificial inhalation, “no-sniff” condition) were measured by calculating spikes / second (Hz) for the 9 sec before or during stimulation. Selection of ‘sniff modulated’ units was performed as described previously ^28^. Inhalation-evoked responses during inhalation of clean air (“sniff” condition) were measured by averaging the number of spikes per 1-sec period following each inhalation in the 9 inhalations pre-stimulation or during stimulation and across multiple trials (minimum of 3 trials in each condition for all units). Odorant-evoked responses were measured as changes in the mean number of spikes evoked per 1-sec inhalation cycle (Δ spikes / sniff) during odorant presentation, relative to the same number of inhalations just prior to odorant presentation. For statistical analysis, significance for changes in firing rate for baseline versus optical stimulation was tested on a unit-by-unit basis using the Mann-Whitney *U* test on units tested with 5 or more trials per condition.

### Statistical analysis

Significance was determined using paired Student’s *t*-test, Wilcoxon signed rank test and Mann– Whitney U test, where appropriate. Significance was defined as *P<0.05, **P<0.005, ***P<0.0005, ****P<0.0001. All tests are clearly stated in the main text.

### Histology

Viral (AAV.FLEX.ChR2.YFP, AAV.FLEX.HR.YFP) expression in BF cells / axonal projections was evaluated with post hoc histology in all experiments to confirm accurate targeting of BF neurons and a lack of expression in OB neurons as described in ^28,104^. Briefly, mice were deeply anesthetized with an overdose of sodium pentobarbital and perfused with 4% paraformaldehyde in PBS. Tissue sections were evaluated from native fluorescence without immunohistochemical amplification with a Leica TCS SP2 confocal laser scanning microscope at 10x or 20x magnification. The fluorescence intensity of confocal images was analyzed and plotted in ImageJ. Only mice showing a solid OB fibre expression were used for analysis.

## Acknowledgments

The authors thank the technical workshop at the institute for biology II, RWTH Aachen for excellent technical support. We thank all lab members for helpful discussion and comments on the manuscript. We thank R. Medinaceli Quintela, L. Wallhorn, V. Schweda J. Koesling and N. Rose for assistance with optogenetics experiments, especially in the early project phase. Initial data were acquired in Utah. We thank Drs. Looger, Akerboom and Kim and the Genetically Encoded Calcium Indicator (GECI) Project at Janelia Farm Research Campus in collaboration with Penn Vector Core for providing with opsin-expressing viruses. This work was funded by the Deutsche Forschungsgemeinschaft (DFG, German Research Foundation) - (RO4046/2-1 and /2-2, Emmy Noether Program [to MR] and the Research Training Group 2416 “MultiSenses – MultiScales: Novel approaches to decipher neural processing in multisensory integration” 368482240/GRK2416).

## Author contributions

EB performed experiments and analyzed data; DB and MR designed experiments, analyzed data and wrote the manuscript

## Additional Information

Competing interests: The authors declare no competing interests.

## Supplementary Information

**Supplemental Figure 1.**
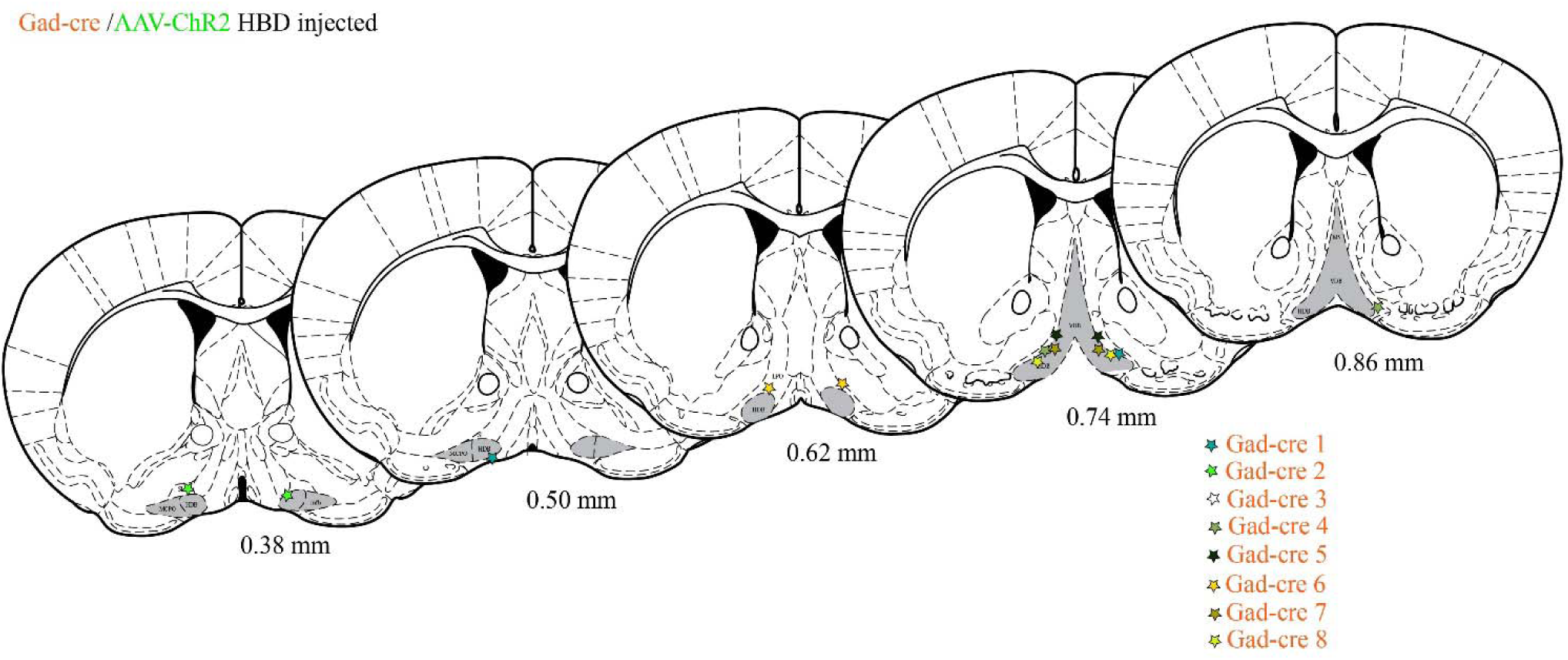
Histological reconstruction of basal forebrain injection sites in GAD-Cre mice. Section from the atlas ^1^ illustrate the reconstructed injection sites from eight GAD-Cre animal injected with AAV-ChR2. HDB/VDB BF area is marked in grey.

**Supplemental Figure 2.**
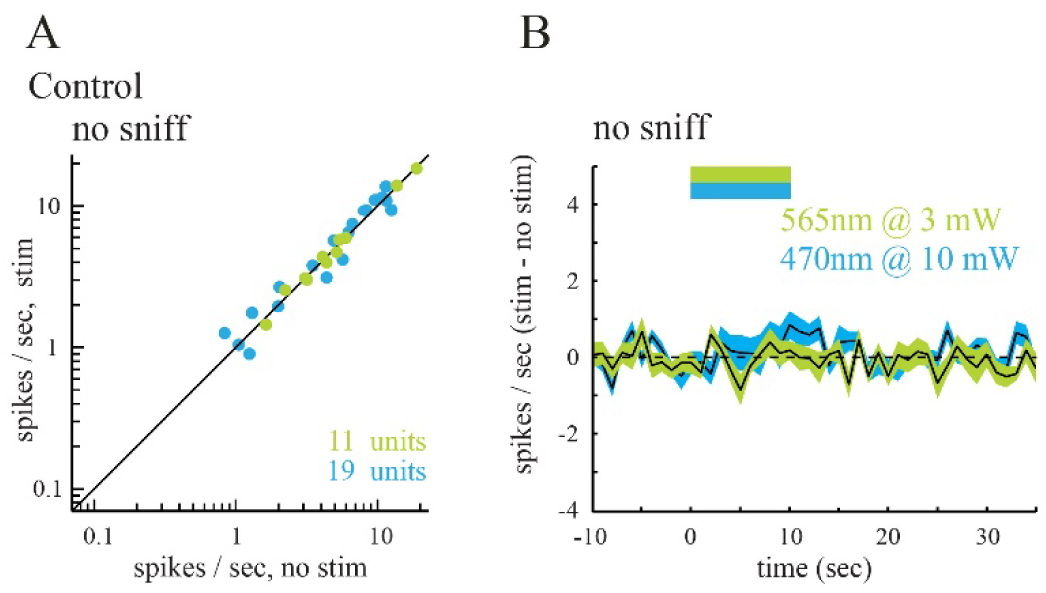
Light-evoked modulation of spontaneous MTC firing in ChAT-Cre and GAD-Cre mice is attributable to cholinergic/GABAergic signaling. A. Plot of spontaneous firing rate of MTCs in control, uninjected ChAT-Cre or GAD-Cre mice before and during optical stimulation of the OB (n = 11 units, 470nm @ 10 mW, blue; n = 19 units 565 nm @ 3 mW, green), recorded and analyzed as in Fig 2B. B. Time course of firing rate change across all recorded MTCs during optical stimulation in control mice.

**Supplemental Figure 3.**
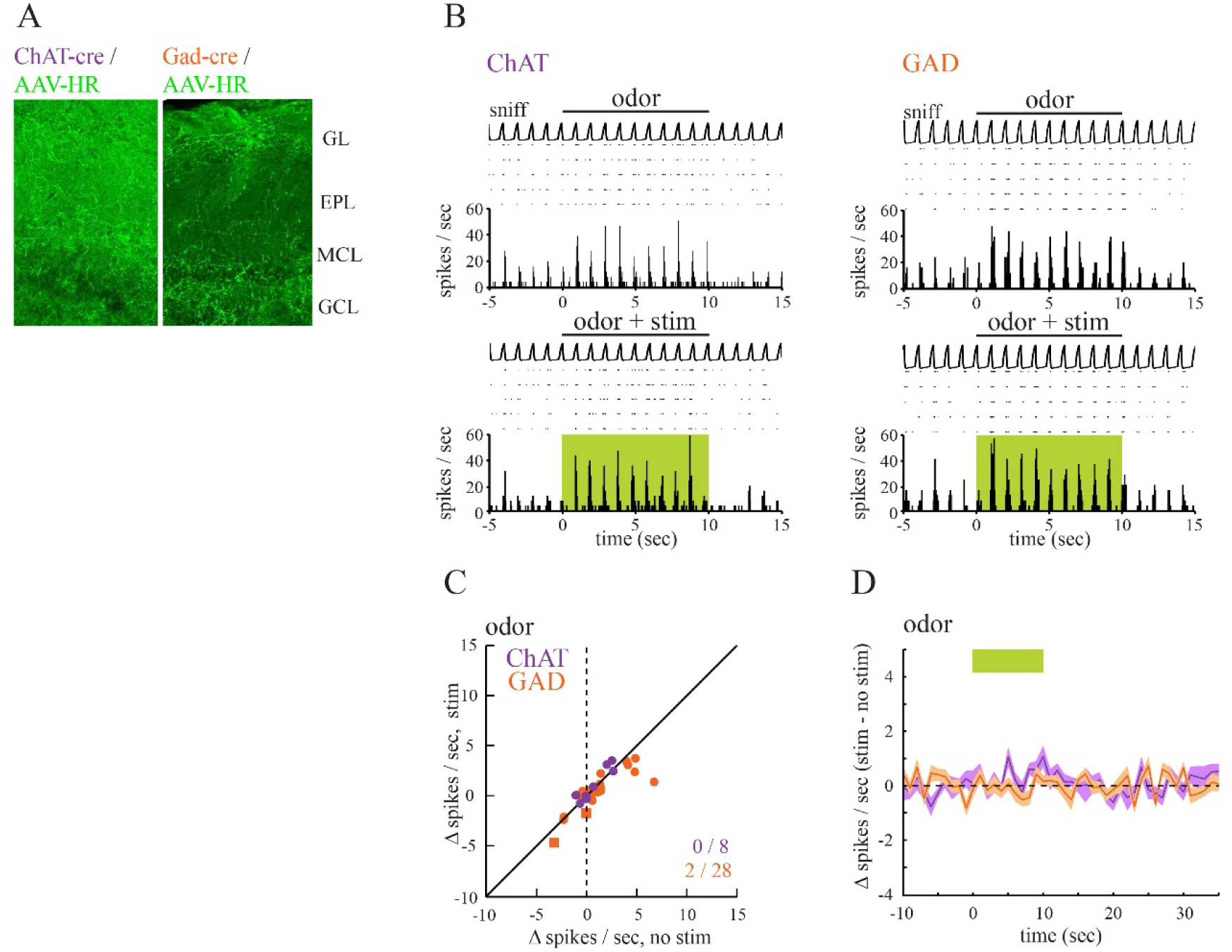
Effects of optogenetic silencing of cholinergic and GABAergic OB inputs on MTC odor-evoked responses. A. HR-EYFP-expressing axon terminals in the OB imaged 4 weeks after AAV-HR-EYFP injection into BF of a ChAT-Cre and GAD-Cre mice. GL: glomerular layer, EPL: external plexiform layer, MCL: mitral and tufted cell layer, GCL: granule cell layer. B. Odorant-evoked MTC spiking is response to optical OB silencing in ChAT-Cre and GAD-Cre mice. C. Plot of odorant-evoked changes in MTC spiking (Δ spikes/sniff) in the absence of (no stim) and during (stim) optogenetic silencing of cholinergic (n = 8 units, purple) or GABAerig (n = 28, orange) afferents to the OB. D. Time course of effects of optical silencing on odorant-evoked spike rate, averaged across all units. The green bar shows time of optical stimulation and simultaneous odorant presentation. The shaded area indicates variance (SEM) around mean.

**Supplemental Figure 4.**
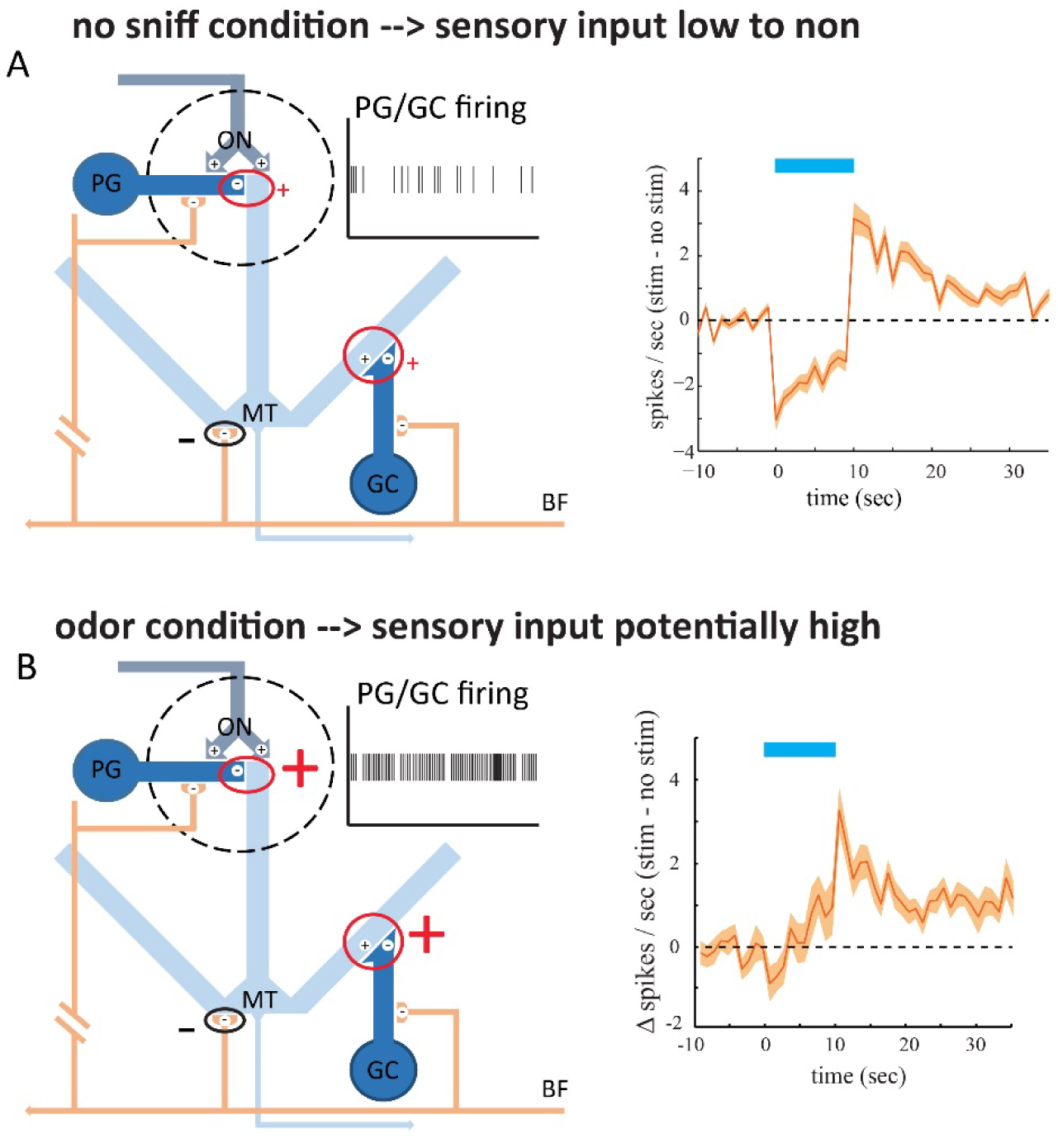
Possible mechanism of input dependent GABAergic modulation of MTCs. Schematic diagram summarizing the connectivity of the GABAergic projection from basal forebrain (adapted from ^2^, GC= granule cells PG= periglomerular cells, ON= olfactory nerve, MT= mitral/tufted cells). A: In the no sniff condition, with little to no sensory input, inhibitory interneurons like PGs and GCs show little activity. This means that GABAergic BF derived inhibition of these cells has little effect and little to none disinhibition of MTCs can be observed. At the same time direct inhibition of BF derived fibers on MTCs might mediate the inhibitory effect seen in this condition (right). B: In the odor condition, with strong sensory input, inhibitory interneurons like PGs and GCs show stronger activity. Therefore, GABAergic BF derived inhibition of these cells can lead to disinhibition of MTCs potentially outweighing the direct inhibition of MTCs.

